# Genetic screening identifies a SUMO protease dynamically maintaining centromeric chromatin and the associated centromere complex

**DOI:** 10.1101/620088

**Authors:** Sreyoshi Mitra, Dani L. Bodor, Ana F. David, João F. Mata, Beate Neumann, Sabine Reither, Christian Tischer, Lars E.T. Jansen

**Author notes:** MRC - Laboratory for Molecular Cell Biology, UCL, London, United Kingdom. Institute of Molecular Biotechnology, Dr. Bohr-Gasse 3, 1030 Vienna, Austria.

## Abstract

Centromeres are defined by a unique self-propagating chromatin structure featuring nucleosomes containing the histone H3 variant CENP-A. CENP-A turns over slower than general chromatin and a key question is whether this unusual stability is intrinsic to CENP-A nucleosomes or rather imposed by external factors. We designed a specific genetic screen to identify proteins involved in CENP-A stability based on SNAP-tag pulse chase labeling. Using a double pulse-labeling approach we simultaneously assay for factors with selective roles in CENP-A chromatin assembly. We discover a series of new proteins involved in CENP-A propagation, including proteins with known roles in DNA replication, repair and chromatin modification and transcription, revealing that a broad set of chromatin regulators impacts in CENP-A transmission through the cell cycle. The key factor we find to strongly affect CENP-A stability is SENP6. This SUMO-protease controls not only the levels of chromatin bound CENP-A but is required for the maintenance of virtually the entire centromere and kinetochore, with the exception of CENP-B. Acute depletion of SENP6 protein reveals its requirement for maintaining centromeric CENP-A levels throughout the cell cycle, suggesting that a dynamic SUMO cycle underlies a continuous surveillance of the centromere complex.

## Introduction

Human centromeres are defined by an unusual chromatin domain that features nucleosomes containing the H3 variant CENP-A (Palmer et al., 1987, 1991; Sekulic and Black, 2012; Black et al., 2010), also known as cenH3. While these nucleosomes typically assemble on α-satellite sequences, they undergo a chromatin-based self-templated duplication along the cell cycle that is largely independent from local DNA sequence features (Mendiburo et al., 2011; Hori et al., 2013; Barnhart et al., 2011). Indeed, CENP-A is sufficient to initiate a centromere and render it heritable through the mitotic cell cycle (Mendiburo et al., 2011; Hori et al., 2013). The maintenance of centromeric chromatin depends on 1) stable transmission of CENP-A nucleosomes across multiple cell cycles (Bodor et al., 2013; Jansen et al., 2007; Falk et al., 2015), 2) a template directed assembly mechanism that depends on previously incorporated CENP-A (Barnhart et al., 2011; French et al., 2017; Hori et al., 2017; Moree et al., 2011; Dambacher et al., 2012; Wang et al., 2014) and 3) cell cycle regulated inheritance and assembly to ensure centromeric chromatin is replicated only once per cell cycle (Jansen et al., 2007; Silva et al., 2012; Stankovic et al., 2017; Schuh et al., 2007; Moree et al., 2011; Bernad et al., 2011; Shukla et al., 2018).

Several factors involved in the assembly of CENP-A histones into centromeric nucleosomes have been identified (Erhardt et al., 2008; Maddox et al., 2007; Fujita et al., 2007; Foltz et al., 2009; Chen et al., 2014; Barnhart et al., 2011). However, comparatively little is known about how CENP-A, once assembled into chromatin, is stably transmitted from one cell cycle to the next. This is relevant as we previously found CENP-A to have a longer half-life than the canonical histone counter parts (Bodor et al., 2013). Photoactivation experiments have shown that this unusually high degree of stability is restricted to centromeres (Falk et al., 2015), a finding recently borne out in genome wide ChIP analysis (Nechemia-Arbely et al., 2018). This suggests that the degree of CENP-A retention in chromatin is a regulated process that is dependent on the centromere. Indeed, an initial insight came from biophysical measurements of the CENP-A nucleosome in complex with its direct binding partner CENP-C that stabilizes CENP-A nucleosomes both in vitro and in vivo (Falk et al., 2015, 2016; Guo et al., 2017) although it is not known whether this occurs at a specific transaction along the cell cycle.

As stable transmission of CENP-A nucleosomes appears central to the epigenetic maintenance of centromeres we sought to identify novel factors that specifically control CENP-A stability in chromatin. Previous genetic screens assayed for factors involved in CENP-A localization (Erhardt et al., 2008; Fu et al., 2019). While those efforts focused on measuring changes in steady state centromere levels of CENP-A, we devised a dedicated assay that allows for the discovery of factors specifically involved in either CENP-A maintenance or CENP-A assembly or both. We have previously employed SNAP tag-based fluorescent pulse-chase labeling to visualize the turnover of chromatin-bound CENP-A molecules across multiple cell divisions (Bodor et al., 2013, 2012). In addition, using an adapted quench-chase-pulse labeling technique we have used SNAP to track the fate of nascent CENP-A (Jansen et al., 2007; Silva et al., 2012; Stankovic et al., 2017; Bodor et al., 2013, 2012). We constructed a custom siRNA library consisting of targets against 1046 different genes encoding chromatin associated proteins and coupled this to a combined pulse-chase and quench-chase-pulse labeling strategy and high throughput imaging. We identified a host of novel proteins, not previously associated with CENP-A metabolism that are involved specifically in either CENP-A loading or maintenance of the chromatin bound pool of CENP-A. We find factors with known roles in DNA replication, repair and chromatin modification and transcription.

The most prominent candidate resulting from our screen is SENP6, a SUMO-specific protease, depletion of which resulted in strong defects in CENP-A maintenance. Upon exploring the broader centromere and kinetochore complex, we find SENP6 to control the localization of all key centromere proteins analyzed, including CENP-C and CENP-T and the downstream kinetochore. Importantly, loss of centromere localization of CENP-A does not affect its cellular levels, excluding a major role for SUMO-mediated polyubiquitination and proteasomal degradation. Finally, using a controllable degron-allele we find that SENP6 is continuously required to maintain the centromeric chromatin throughout the cell cycle.

## Results

### Design of a genetic screening platform to identify specific CENP-A assembly and CENP-A maintenance factors

We built upon our previous expertise to design a SNAP tag-based screening strategy to identify factors involved in CENP-A maintenance as well as in the assembly of new CENP-A. SNAP-pulse labeling allows for the differential labeling of either the pre-existing protein pool or of the newly synthesized pool (Figure 1A). By using two spectrally distinct fluorophores we can track both the old pre-incorporated centromeric CENP-A pool over the course of several cell divisions and the new CENP-A pool simultaneously within the same cells and at the same centromeres (Figure 1B).

**Figure 1.**
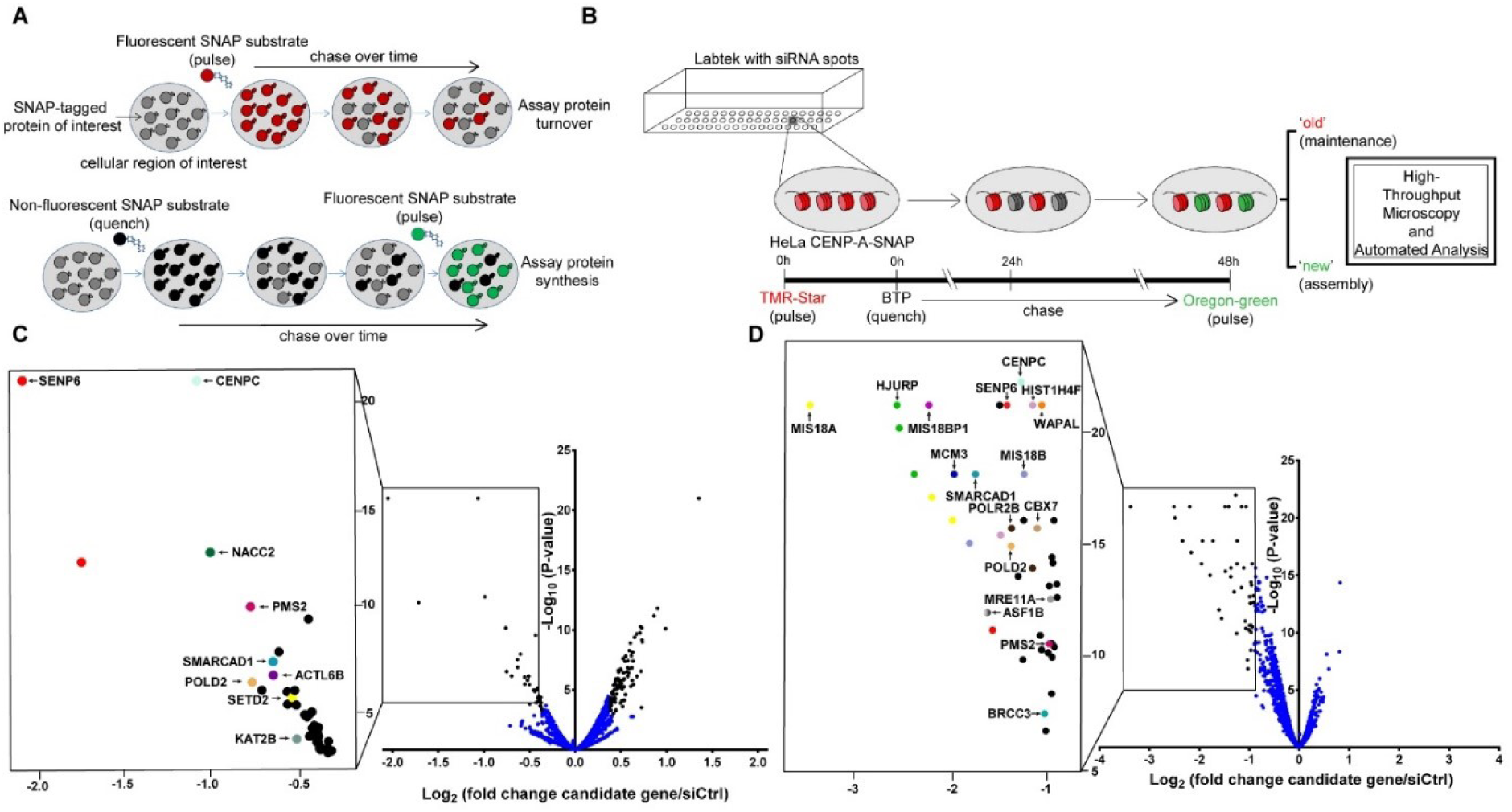
An imaging-based genetic screen identifies potential regulators of CENP-A maintenance and assembly. **(A)** Schematic depicting the SNAP tag-based labeling assay in order to track the maintenance and turnover (red) or de novo assembly (green) of a SNAP-tagged protein. **(B)** Flow scheme of high throughput siRNA screen designed to identify novel candidates controlling the loading or maintenance of CENP-A. HeLa CENP-A-SNAP cells, pulse labelled with TMR-Star were seeded onto chamber slides carrying 384 siRNA printed spots. During RNAi, cells were chased for 48 hours followed by labeling of nascent CENP-A-SNAP with Oregon Green. CENP-A maintenance or assembly was assessed by high throughput fluorescence microscopy of red (old) or green (new) CENP-A-SNAP signals, respectively. **(C)** Volcano plot representing the results of the siRNA screen scoring for defects in maintenance of pre-assembled CENP-A. The fold change of mean ‘old’ CENP-A intensity in the candidate siRNA treatment versus that of a negative scrambled siRNA control are plotted on the x-axis in log_2_ scale. The p-values of the candidate siRNA treatment as a measure of reproducibility across 5 biological replicates of the screen are plotted on the y-axis in negative log_10_ scale. The top candidate maintenance factors with a cut-off lower than a fold change (log_2_) of −0.4 and higher than a p-value (−log_10_) threshold of 3 are boxed and highlighted (identically colored dots represent different siRNA targets for the same gene). **(D)** Volcano plot as in (C) representing the results of the siRNA screen scoring for defects in CENP-A assembly. The top candidate assembly factors with a cut-off lower than a fold change (log_2_) of −0.9 and higher than a p-value (−log_10_) threshold of 5 are boxed and highlighted.

We assembled a custom made siRNA library representing genes involved in a broad set of chromatin functions, including DNA replication, transcription, DNA repair, SUMO and Ubiquitin-regulation, nuclear organization, chromatin remodeling and histone modifications. Our library is comprised of 2172 siRNAs encompassing 1046 genes (see supplementary table S1).

To screen for the involvement of these genes in CENP-A maintenance and/or assembly, we spotted 4 nanoliters of an siRNA/Lipofectamine mixture onto chambered cover glass using contact printing technology. A HeLa cell line expressing near endogenous levels of CENP-A-SNAP (Jansen et al., 2007) was SNAP pulse labeled with a rhodamine-conjugated SNAP substrate for chromatin bound CENP-A and was solid-reverse transfected by seeding onto slides carrying 384 siRNA spots each. Following the initial pulse labeling and siRNA-mediated target mRNA depletion, cells were allowed to undergo 2 rounds of cell divisions and CENP-A turnover after which nascent CENP-A was pulse labeled using an Oregon green-conjugated SNAP substrate. Centromeric fluorescence intensity was determined for each siRNA condition using an automated centromere imaging pipeline (Figure 1B, Figure S1, see methods for details).

### Genetic screening identifies novel CENP-A assembly and maintenance factors

We screened all 2172 siRNAs in 5 replicate experiments which allowed us to determine both the difference in CENP-A intensity for each siRNA relative to a scrambled control siRNA as well as the variance in these differences. We identified 33 siRNAs (representing 31 genes) that led to a reduced CENP-A maintenance above a threshold of 1.3 fold (−0.4 on Log_2_ scale) relative to controls with a significance higher than p = 0.001 (3 on -Log_10_ scale) (Figure 1C and Table 1, supplementary Table S2). Among the putative genes whose depletion has a significant impact is CENP-C, which was previously found to stabilize CENP-A (Falk et al., 2015) and served as a control in our screen. In addition we find groups of genes involved particularly in **1) SUMO/Ubiquitin** transactions, such as SENP6, a SUMO protease (Mukhopadhyay and Dasso, 2007), **2) chromatin remodelers** that include SMARCAD1, a SWI/SNF type chromatin remodeler involved in chromatin reconstitution following DNA replication (Taneja et al., 2017; Mermoud et al., 2011), **3) chromatin modifiers such as** SETD2 a H3K36 histone methyltransferase (Edmunds et al., 2008; Yuan et al., 2009), **4)** factors involved in **transcription regulation**, notably NACC2, a POZ domain containing protein (Xuan et al., 2013) and also **5) DNA replication and repair factors** such as the DNA mismatch repair protein PMS2 (Kolodner and Marsischky, 1999). See pathway-clustered groups of genes in table 1.

**Table 1.**
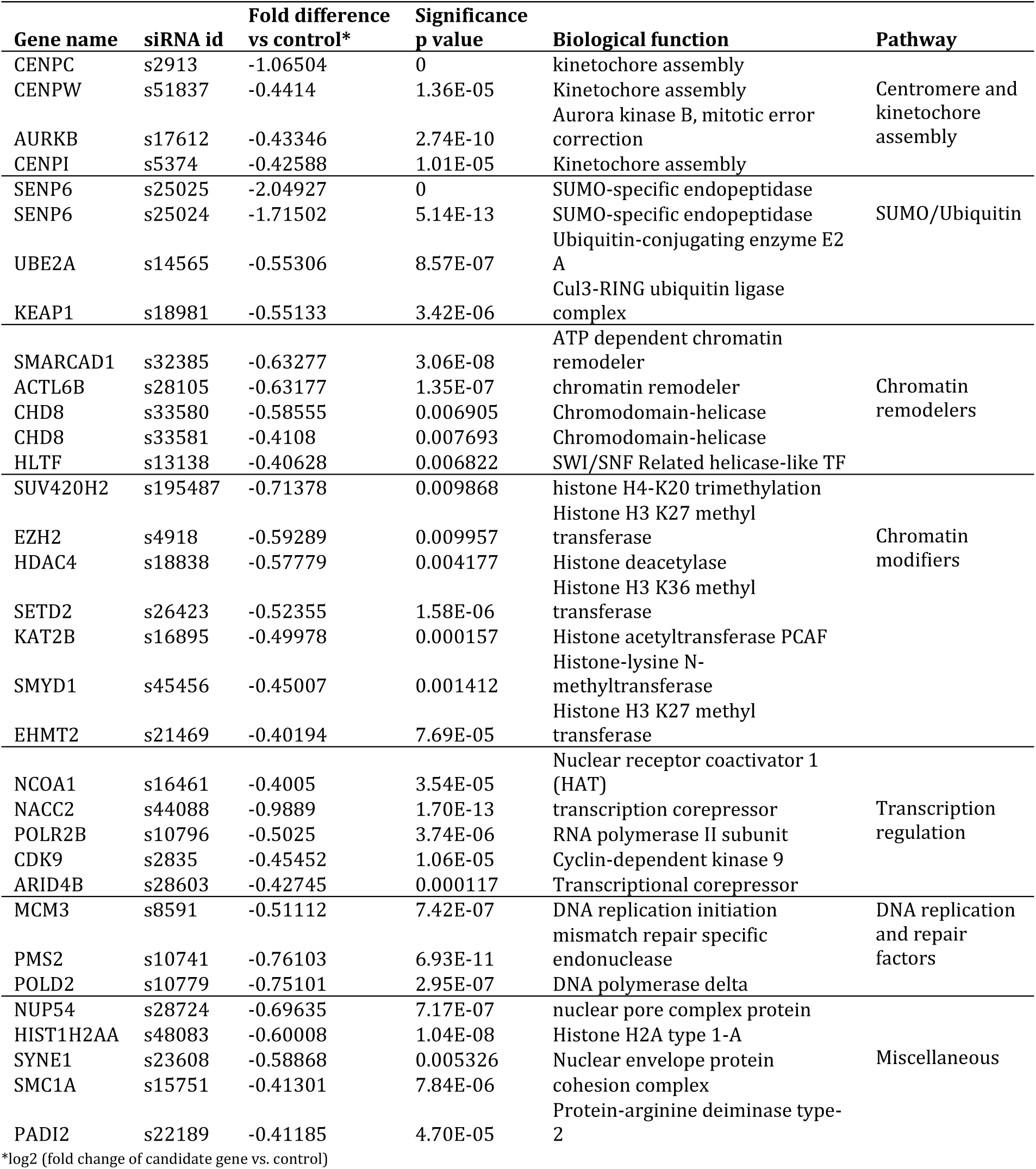
List of candidate genes that affect the maintenance of ‘old’ CENP-A molecules at the centromere. List of candidate genes affecting maintenance of pre-assembled centromeric CENP-A, clustered in pathways. Listed are hits with a fold difference higher than 1.3 (−0.4 on log 2 scale) of mean ‘old’ CENP-A signal intensity in the siRNA treatment versus the negative scrambled siRNA control and have a significance over <0.001.

The same 2172 siRNA panel revealed a distinct set of genes when screening for factors involved in loading of nascent CENP-A (Figure 1D and Table 2, supplementary Table S3). We identified 42 siRNAs (representing 34 genes) that led to a reduced CENP-A assembly above a threshold of 1.9 fold (−0.9 on Log_2_ scale) relative to controls with a significance higher than p = 0.0001 (5 on -Log_10_ scale) (Figure 1C and Table 2, supplementary Table S3). Among these we found known assembly factors including the CENP-A specific chaperone HJURP and all members of the Mis18 complex, M18BP1, Mis18α and M18β as well as CENP-C. Further we identified several novel components not previously associated with CENP-A assembly. Among DNA replication and repair factors we find MCM3 (Alabert and Groth, 2012) and POLD2 as well as the chromatin remodeler SMARCAD1 (Mermoud et al., 2011) and the transcription regulator CBX7, a Polycomb repressive complex 1 (PRC1) member (Blackledge et al., 2015). See pathway-clustered groups of genes in tables 1, 2, S2 and S3 and further description of the candidate factors in the discussion.

**Table 2.**
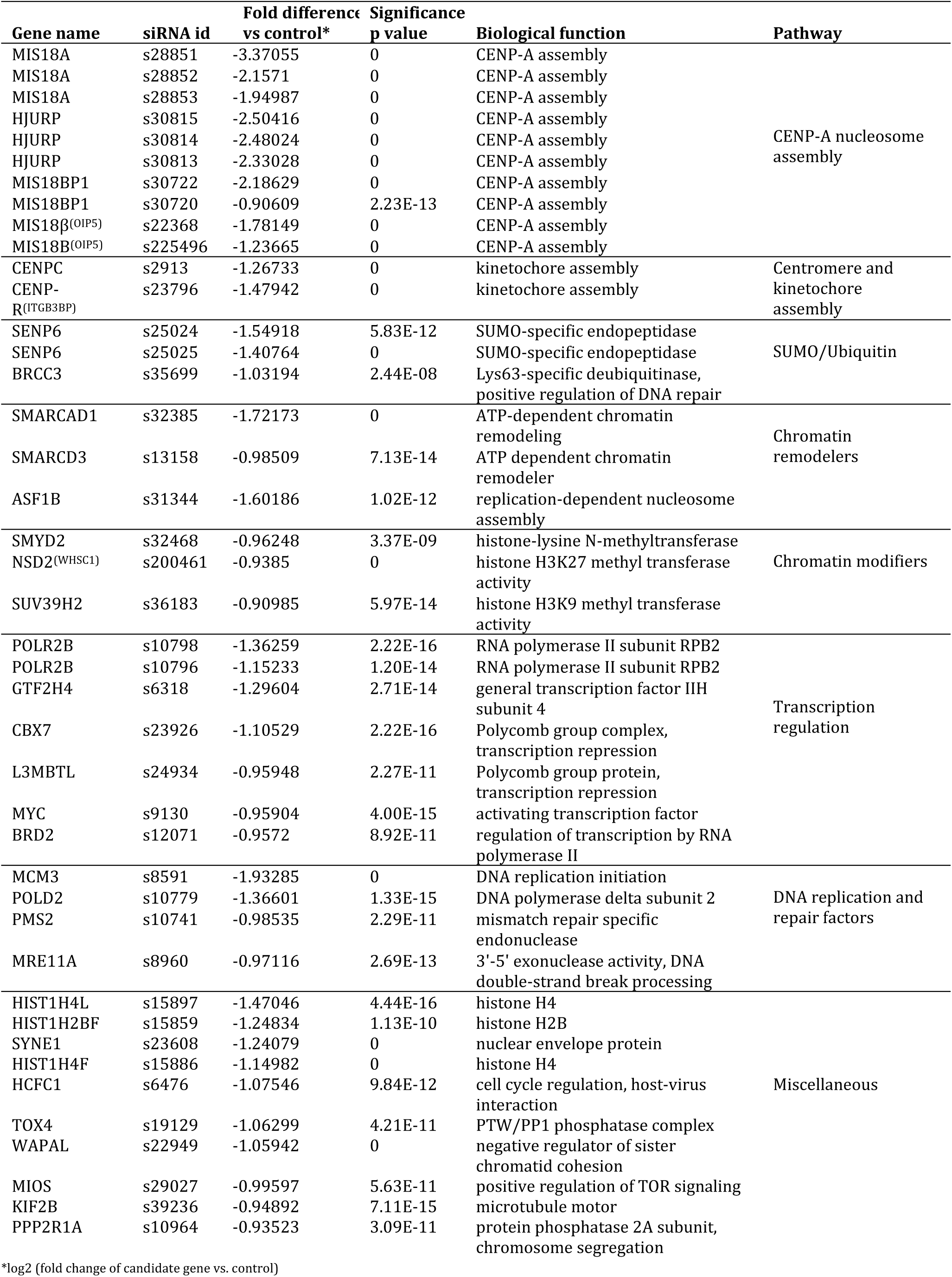
List of candidate genes that affect the loading of ‘new’ CENP-A molecules at the centromere. List of candidate genes affecting assembly of nascent CENP-A at centromeres, clustered in pathways. Listed are hits with a fold difference higher than 1.9 (−0.9 on log 2 scale) of mean ‘new’ CENP-A signal intensity in the siRNA treatment versus the negative scrambled siRNA control and have a significance over <0.0001.

Strikingly, the top two siRNAs reducing CENP-A maintenance are both against SENP6 (4.1 and 3.3 fold reduction,2.05 and 1.72 on Log2 scale (Table 1)). SENP6 also scored high as a factor affecting CENP-A assembly (Table 2) suggesting that SENP6 is required for maintaining all CENP-A nucleosomes, including recently incorporated CENP-A. Alternatively, it has an additional role specifically in assembly. The importance of SENP6 in CENP-A turnover is further highlighted by a recent independent study, where it was identified as a factor involved in CENP-A centromere localization (Fu et al., 2019). This study implicated M18BP1 as a mediator of SENP6 imposed CENP-A stabilization. However, we do not find M18BP1 to have a significant impact on CENP-A maintenance, while playing a major role in depositing new CENP-A in the same cells in our double labeling setup (Figure 1D and supplementary Table S2, S3), consistent with previous results (Fujita et al., 2007; Bodor et al., 2013; Dambacher et al., 2012; Moree et al., 2011).

### SENP6 controls CENP-A maintenance

The SUMO protease SENP6 is involved in proteolytic removal of SUMO2/3 chains from target proteins (Mukhopadhyay and Dasso, 2007). SENP6 is known to be involved in various processes including inflammatory signaling and DNA repair (Dou et al., 2010; Liu et al., 2013). Furthermore, SENP6 has previously been implicated in regulating the levels of CENP-H and CENP-I at the centromere (Mukhopadhyay et al., 2010). To explore the involvement of SENP6 in CENP-A regulation we depleted SENP6 from HeLa CENP-A-SNAP cells by siRNA. We confirmed at high resolution and single centromere intensity measurements that SENP6 affects maintenance of both the CENP-A pool that was incorporated into chromatin prior to SENP6 depletion (Figure 2A, C, E) as we all as maintenance of the nascent CENP-A pool (Figure 2B, D, E). As expected, centromeric steady state levels of CENP-A were also reduced (Figure 2G, H).

**Figure 2.**
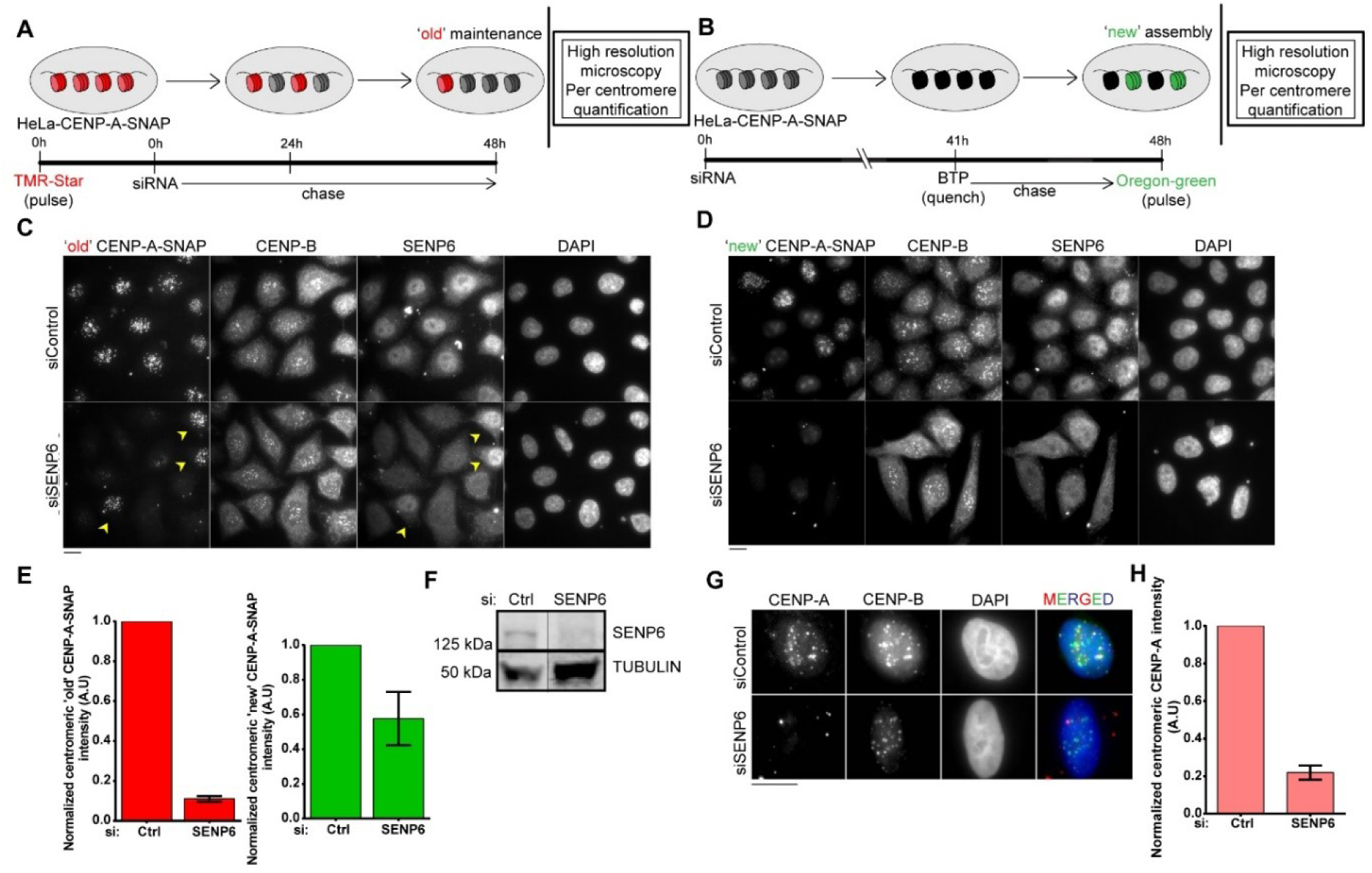
SENP6, a SUMO protease, controls maintenance of centromeric CENP-A. **(A)** and **(B)** Schematics for high resolution SNAP pulse-chase (A) and quench-chase-pulse (B) assays. HeLa-CENP-A-SNAP cells were treated with SENP6 siRNA or a control scrambled siRNA. Pulse-chase experiment was performed for 48 hours during RNAi to assay for CENP-A turnover (A). Quench-chase-pulse experiment was performed for the final 7 hours of siRNA treatment to assay for CENP-A assembly (B). **(C)** and **(D)** shows typical image fields following the strategies in (A) and (B) respectively. TMR-Star and Oregon Green SNAP labels visualize the maintenance or assembly of CENP-A-SNAP, respectively. CENP-B was used as a centromeric reference for quantification. Cells were counterstained for SENP6 to visualize its depletion in siRNA treated cells. Yellow arrowheads indicate nuclei that escaped SENP6 depletion which correlate with retention of ‘old’ CENP-A-SNAP. Bars, 10 µm. **(E)** Automated centromere recognition and quantification of (C) and (D). Centromeric CENP-A-SNAP signal intensities were normalized to the control siRNA treated condition in each experiment. siRNA treatment; siSENP6 or scrambled (Ctrl). Three replicate experiments were performed. Bars indicate SEM. **(F)** Western blot indicating depletion of SENP6 protein following siRNA. Extracts made from SENP6 siRNA treated or control siRNA treated cells were separated by SDS-PAGE and immunoblotted with anti-SENP6 antibody. Tubulin was used as loading control. **(G)** Centromeric levels of total, steady state CENP-A under siRNA mediated depletion of SENP6. **(H)** Quantification of (G). Three replicate experiments were performed. Bars indicate SEM.

### SENP6 is involved in maintaining the entire centromere complex and associated kinetochore independent of proteolysis

Next, we determined whether other centromere components were also affected by depletion of SENP6. Previous reports have indicated a requirement of SENP6 for maintaining CENP-H and –I at centromeres (Mukhopadhyay et al., 2010). We now found that all principle constitutive centromere components CENP-C, CENP-T in addition to CENP-A are lost after depletion of SENP6 (Figure 3A, B, D). These findings indicate that the entire DNA-associated foundation of the centromere is under the critical control of SENP6. A notable exception is CENP-B, an alpha-satellite binding centromere protein, which remains unaffected by SENP6 depletion (Figure 3B, D). Furthermore, key members of the microtubule binding KMN-network, Nnf-1, Dsn-1, and Hec-1 are lost after SENP6 depletion as well (Figure C, D).

**Figure 3.**
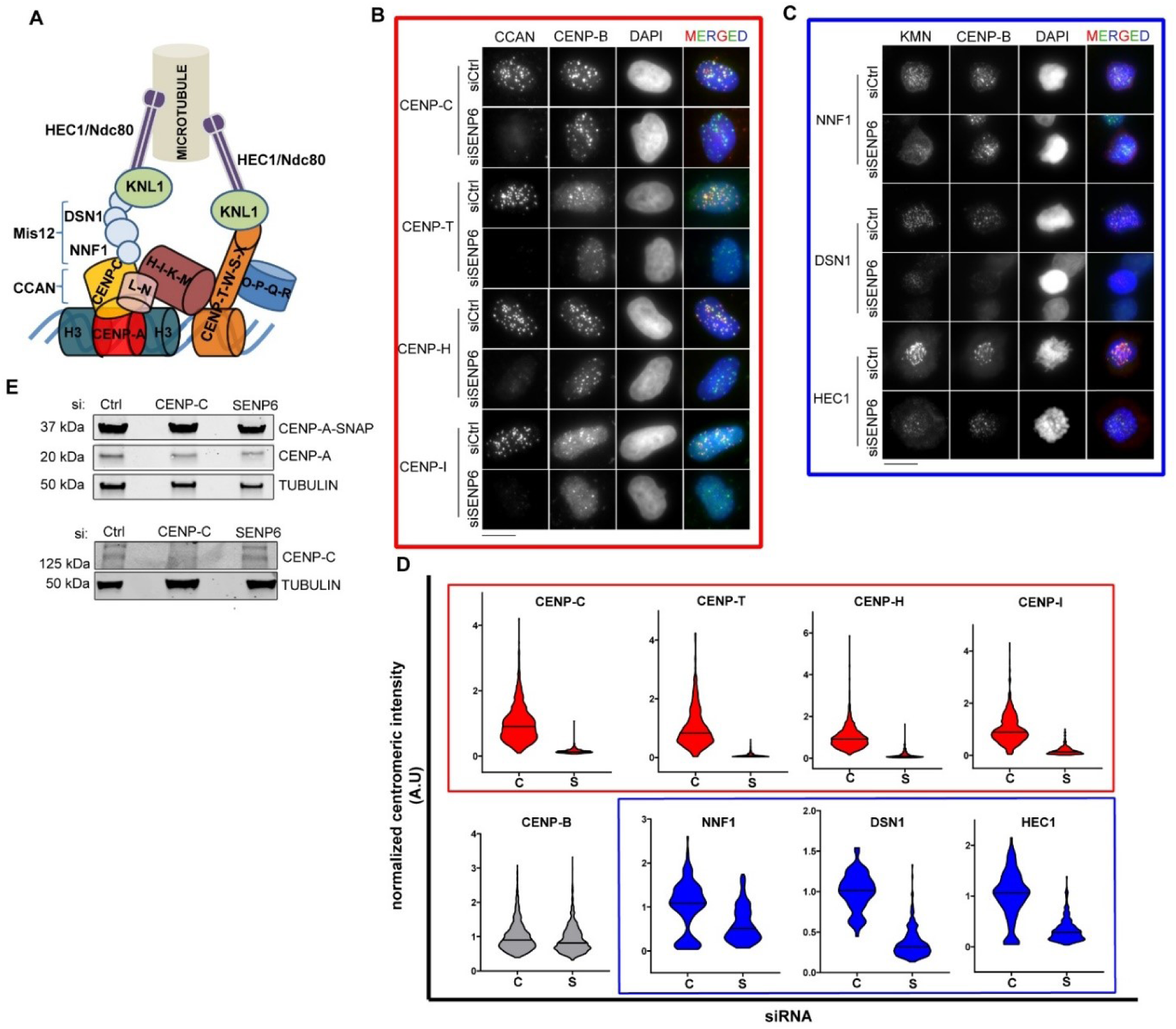
SENP6 is required for maintaining the integrity of the constitutive centromere-associated network (CCAN) and associated kinetochore, independent of proteolysis. **(A)** Schematic representing the architecture and interactions of different protein complexes in the human centromere and kinetochore. **(B)** Centromeric levels of different CCAN proteins following siRNA mediated depletion of SENP6. Cells were counterstained with CENP-B to mark centromeres. **(C)** Centromeric levels of KMN (KNL1-MIS12-NDC80) members following siRNA mediated depletion of SENP6. Bars, 10 µm. **(D)** Automated centromere recognition and quantification of (B) and (C). Fluorescence intensities of indicated proteins were normalized to the mean of the control siRNA treated condition in each experiment and plotted as a violin plot following siRNA of SENP6 “S” or control “C”. At least 200 centromeres were measured for each protein and each treatment. Bar indicates the median value. **(E)** Western blot showing the total levels of CENP-A and CENP-C proteins following 48 hour treatment with siRNAs against SENP6 and CENP-C. Extracts from SENP6 or CENP-C siRNA treated or control siRNA treated cells were separated by SDS-PAGE and immunoblotted with anti-CENP-A (detecting both SNAP tagged and endogenous CENP-A) or anti-CENP-C antibody. Tubulin was used as loading control.

It has been shown that polySUMOylation can lead to subsequent polyubiquitination by SUMO-directed ubiquitin ligases such as RNF4. This mechanism was previously implicated in the degradation of CENP-I (Mukhopadhyay et al., 2010). As a deSUMOylase, SENP6 could prevent CENP-I, and other centromere/kinetochore protein, degradation by removing SUMO chains. Indeed, SUMO-mediated polyubiquitination and subsequent degradation of budding yeast CENP-A^Cse4^ has been reported (Ohkuni et al., 2018, 2016). Therefore, we measured the protein stability of CENP-A and -C after depletion of SENP6. Unexpectedly, we find that while these proteins are lost from centromeres, cellular levels of both proteins are maintained upon SENP6 depletion (Figure 3E). This suggests that hyper-SUMOlyation of target proteins in SENP6 depleted cells does not lead to polyubiquitination and proteasomal degradation but rather that a SUMO cycle controls localization.

### SENP6 is continuously required to maintain CENP-A at the centromere

A critical barrier for chromatin maintenance is S phase during which DNA replication disrupts histone DNA contacts. A key step in the stable propagation of chromatin is histone recycling during DNA replication involving the MCM2-7 helicase complex, along with dedicated histone chaperones (Huang et al., 2015), a role recently extended to the stable transmission of CENP-A nucleosomes at the replication fork (Zasadzińska et al., 2018). Further, recent work has shown that the CENP-A chaperone, HJURP is required to recycle CENP-A specifically during S phase (Zasadzińska et al., 2018). In our screen, HJURP is a not a prominent contributor to overall CENP-A maintenance, relative to SENP6 or other candidate maintenance factors (Supplemental Table S2). Nevertheless, S phase may be a critical cell cycle window where CENP-A maintenance requires the re-assembly of pre-existing CENP-A chromatin onto nascent DNA. To determine whether SENP6 plays a role in CENP-A recycling and maintenance of centromere integrity during DNA replication or has broader roles along the cell cycle we constructed a SENP6 specific auxin-inducible degron (AID) (Nishimura et al., 2009) that has been successfully exploited previously to address the acute role of centromere proteins (Holland et al., 2012; Hoffmann et al., 2016; Fachinetti et al., 2015; Guo et al., 2017; Zasadzińska et al., 2018).

We targeted all SENP6 alleles in HeLa cells, which express the CENP-A-SNAP transgene as well as the *O. sativa*-derived E3 ligase, TIR1 with a miniAID-GFP construct (Figure 4A, S2A). Addition of the auxin Indole-3-acetic acid (IAA) resulted in rapid loss of SENP6 which became undetectable upon a 3 hour incubation (Supplemental Figure S2A, B). Longer exposure to IAA resulted in cell growth arrest confirming SENP6 to be an essential protein for cell viability (Supplemental Figure S2C). In agreement with the siRNA experiments above, SENP6 degradation over a 48 hour period let to a loss of CENP-A from centromeres in SNAP-based pulse-chase measurements (Figure 4B, C). Similar to siRNA depletion of SENP6, acute loss of centromeric CENP-A after SENP6 protein degradation did not lead to a loss in cellular levels of CENP-A or CENP-C (Supplemental Figure S2E), suggesting delocalization does not lead to-or is a consequence of degradation. Strikingly, time course experiments of IAA addition showed that loss of CENP-A becomes evident within 6 hours of SENP6 depletion (Figure 4D). The acute effect of SENP6 depletion on CENP-A nucleosomes enables us to determine at what stage during the cell cycle, CENP-A stability depends on SENP6 action.

**Figure 4.**
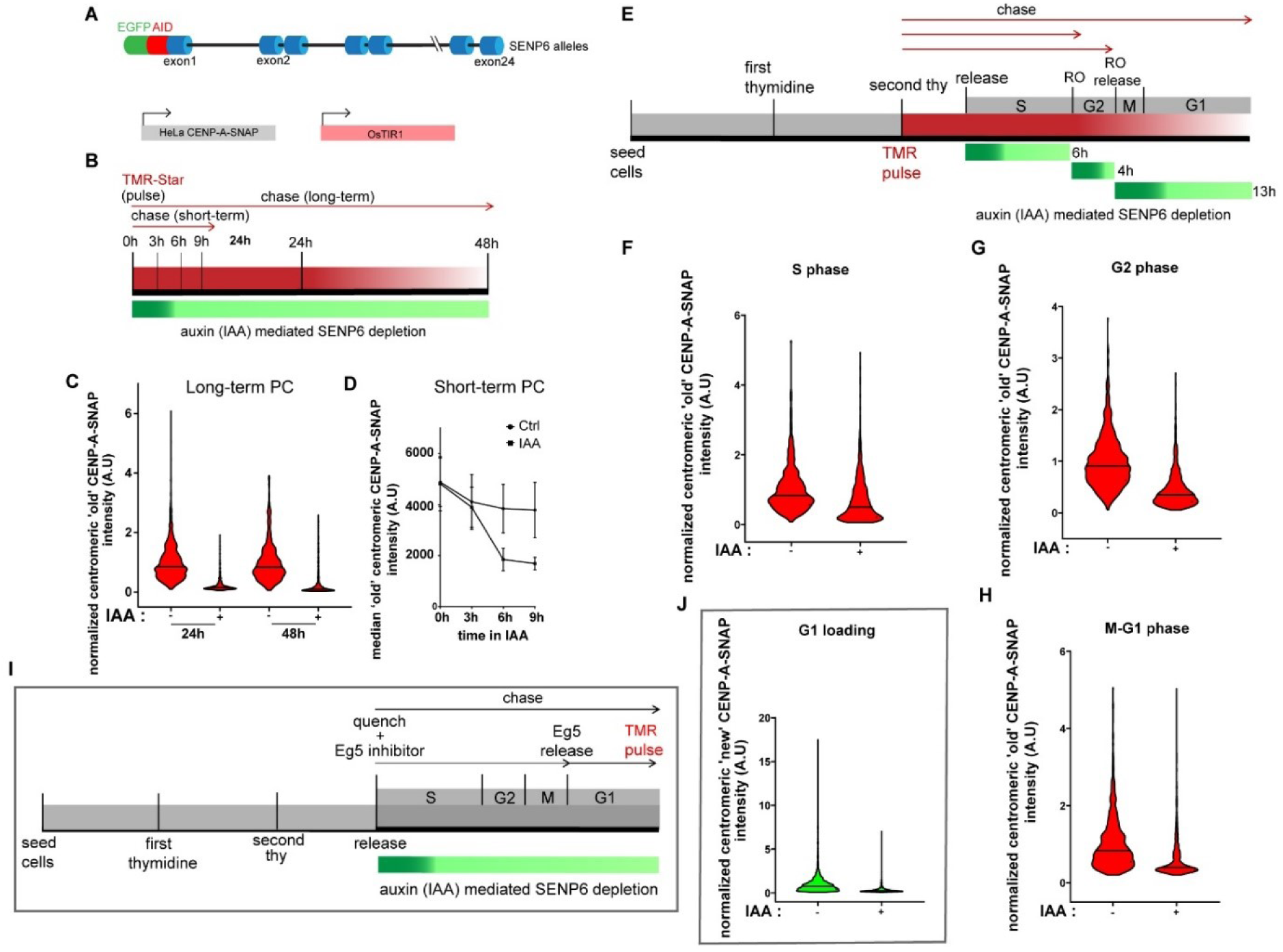
SENP6 is required for centromeric CENP-A maintenance throughout the cell cycle. **(A)** Schematic of the genotype of cell line constructed for auxin (IAA)-mediated depletion of SENP6. OsTIR1 and CENP-A-SNAP are expressed as transgenes, SENP6 is homozygously tagged at its endogenous locus. **(B)** Experimental scheme for long-and short-term CENP-A-SNAP pulse-chase (PC) assays following auxin (IAA) mediated depletion of SENP6. **(C)** and **(D)** Quantification of long-term and short-term PC experiments, respectively following the experimental scheme detailed in (B). **(C)** ‘Old’ centromeric CENP-A-SNAP intensities are normalized to the mean of the non-treated condition (−) for the indicated time point and plotted as a violin plot against auxin (IAA) treated (+) and non-treated (−) conditions for 24h and 48h. At least 500 centromeres were measured in each condition. Bar indicates the median value. **(D)** ‘Old’ centromeric CENP-A-SNAP intensities are measured and median intensities are plotted against auxin (IAA) or non-treated (Ctrl) conditions. Bars indicate SEM of three replicate experiments. **(E)** Experimental scheme of cell cycle synchronization coupled to CENP-A-SNAP pulse-chase (TMR pulse) and auxin (IAA) mediated depletion of SENP6 in different stages of the cell cycle (RO: RO3306, Cdk1 inhibitor). **(F)**,**(G)** and **(H)** Quantification as in (C) of ‘old’ CENP-A-SNAP intensities after auxin (IAA) mediated depletion of SENP6 at different stages of the cell cycle following the experimental scheme of (E). **(I)** Experimental scheme of cell cycle synchronization in G1 phase coupled to CENP-A-SNAP quench-chase-pulse and auxin (IAA) mediated depletion of SENP6. EG5 inhibitor Dimethylenastron was used for a mitotic arrest followed by release into G1 **(J)** Quantification as in (C) of ‘new’ CENP-A-SNAP intensities in G1 stage of the cell cycle after auxin (IAA) mediated depletion of SENP6 following the experimental scheme in (I).

To this end, we synchronized HeLa CENP-A-SNAP expressing cells at the G1/S boundary using Thymidine, pulse labeled CENP-A-SNAP followed by release into S phase (Figure 4E). We depleted SENP6 by the addition of IAA for 4 hours specifically during S phase or after entry into G2 phase during an arrest with the Cdk1 inhibitor for 4 hours. To assess the contribution of SENP6 to CENP-A maintenance in G1 phase we synchronized cells in G2 phase with the Cdk1 inhibitor RO3306 and depleted SENP6 following the release from the inhibitor in mitosis and G1. Irrespective of whether SENP6 was depleted in S, G2 or G1 phase, CENP-A was lost from the centromere (Figure 4F-H). This is a striking observation and indicates that CENP-A chromatin is under continued surveillance and at risk of loss, presumably by SUMO modification. It highlights that CENP-A turnover is a process not only coupled to DNA replication but to other cellular processes as well that may include transcription or other chromatin remodeling activities. Similarly, when cells were synchronized in G1 phase by mitotic arrest and release (Figure 4I) and newly synthesized CENP-A was labeled we found that freshly assembled CENP-A cannot be maintained in the absence of SENP6 (Figure 4J). While SENP6 is likely responsible for the maintenance of newly loaded CENP-A our experiments cannot rule out a direct role for SENP6 in CENP-A assembly.

We confirmed these results in unsynchronized cells in which we depleted SENP6 for 7 hours and analyzed CENP-A loss at specific stages of the cell cycle (Figure S3). We scored G1 phase cells by the presence of α-tubulin positive midbodies, S phase cells by EdU pulse labeling to mark for active DNA replication, and G2 phase cells by cytosolic cylin B staining. SENP6 depletion resulted in loss of CENP-A irrespective of cell cycle position. These results indicate that SENP6 is continuously required throughout the cell cycle to prevent CENP-A from being removed from the centromere.

## Discussion

### Novel pathways involved in the assembly and maintenance of centromeric chromatin

The CENP-A nucleosome is a key determinant of the heritable maintenance of centromeres. A key question that remains is how is CENP-A assembled into chromatin and perhaps more enigmatically, how is it stably transmitted across multiple cell division cycles, a property central to the epigenetic propagation of the centromere. To what extent is stable transmission an intrinsic property encoded within the CENP-A histone or is stable chromatin binding imposed by external factors? To gain insight, we devised a comprehensive screen that is specifically designed to identify nuclear chromatin-associated proteins required for maintaining the chromatin-bound pool of CENP-A. We identified a series of putative factors not previously associated with CENP-A dynamics and centromeric chromatin maintenance.

### Common themes among factors involved in CENP-A maintenance

Among the top ranking candidates we find several clusters of functionalities involved in CENP-A chromatin maintenance. Factors that stand out are proteins involved in **centromere and kinetochore function** that includes CENP-C as expected (Falk et al., 2016, 2015) but also CENP-W and CENP-I as well as the mitotic kinase Aurora B, indicating that several layers within the centromere and kinetochore impact on centromeric chromatin maintenance.

In addition, **chromatin remodeling factors or members of ATP dependent motors** are among the list which includes SMARCAD1, part of the SNF subfamily of helicase proteins, which has been implicated in nucleosome turnover at sites of repair (Terui et al., 2018). We find both previously incorporated as well as new CENP-A to be affected by loss of SMARCAD1. The related protein SMARCD3^BAF60C^ (Debril et al., 2004) was identified as well to specifically affect the nascent pool of CENP-A, possibly implicating it in CENP-A assembly. Interestingly, the fission yeast homolog of SMARCAD1, fft3 has been shown to be involved in faithful maintenance of heterochromatin (Taneja et al., 2017) by reducing histone turnover. Possibly it plays an analogous function in CENP-A chromatin maintenance. Other chromatin remodelers include ACTL6B^BAF53B^, an actin-related protein that is a subunit of the BAF (BRG1/brm-associated factor) complex (Olave et al., 2002) in mammals, which is functionally related to SWI/SNF chromatin remodeler complexes (Alfert et al., 2019). HLTF^SMARCA3^, another SWI/SNF family member (Mansharamani et al., 2001) and CHD8 (Chromodomain Helicase DNA Binding Protein 8) (Manning and Yusufzai, 2017) also have a significant impact on CENP-A maintenance.

Related to the chromatin remodelers are factors involved in **transcriptional repression**, NACC2 (also known as RBB) which contains a POZ domain and recruits the NuRD (Nucleosome Remodeling Deacetylase) complex for gene silencing (Xuan et al., 2013), and ARID4B (Wu et al., 2006) a subunit of the histone deacetylase-dependent SIN3A transcriptional corepressor complex. General components of the transcription machinery such as the POLR2B, a subunit of RNA polymerase II and CDK9^P-TEFb^, a critical kinase in the transcription elongation complex (Cho et al., 2010) also impact on CENP-A maintenance as well as assembly (for POLR2B). Pleiotropic effects of depleting these general transcription-related proteins cannot be excluded but nevertheless these factors are of interest as a direct role for transcription in CENP-A maintenance and assembly has been suggested (Bobkov et al., 2019, 2018; Bergmann et al., 2011).

Furthermore, in addition to core transcription components and chromatin remodelers we identified a series of **chromatin modifying enzymes** including SUV420H2, a histone H4K20 methyltransferase (Schotta et al., 2004), EZH2, the PRC2 complex component responsible for H3K27 methylation (Margueron and Reinberg, 2011), SETD2 the principle H3K36 methyl transferase (Edmunds et al., 2008; Yuan et al., 2009), EHMT2 (G9A) is a H3K9 specific methyltransferase (Krishnan et al., 2011) and SMYD1 an SET domain protein that potentially targets histones (Tracy et al., 2018). The variety of modifiers that impact on CENP-A maintenance suggests that a chromatin imbalance, whether it is activating or repressing for gene expression, has a deleterious effect on CENP-A chromatin maintenance. SUV420H2 is of particular interest as histone H4 in the context of CENP-A nucleosomes has been shown to monomethylated at lysine 20 and impact on centromere structure (Hori et al., 2014). It would be of interest to determine whether SUV420H2 is responsible for this modification and can affect CENP-A nucleosome stability directly. In addition, we identified two acetyl transferases with a potential role in maintaining CENP-A levels. These are NCOA1 and KAT2B [also known as PCAF, a component of the p300 complex (Voss and Thomas, 2018)] as well as an deacetylase, HDAC4.

### DNA repair and replication factors affect both CENP-A maintenance and assembly

Our screening setup allows us to differentiate between proteins involved in recruiting CENP-A to the centromere and those that are required to maintain CENP-A in chromatin, once incorporated. However some components that we identified play a role in both. Possibly, this reflects a requirement of those factors in CENP-A maintenance, both CENP-A that was previously incorporated as well as newly incorporated CENP-A. Factors in this category include CENP-C as has been reported before (Falk et al., 2015) but also the DNA replication factors POLD2 (a DNA polymerase subunit), MCM3 [part of the MCM2-7 DNA helicase complex (Alabert and Groth, 2012)] as well as the mismatch repair factor PMS2 that is a component of the MutLα complex (Kolodner and Marsischky, 1999). Interestingly, MCM2, another member of the MCM2-7 complex has recently been implicated in CENP-A maintenance during S phase (Zasadzińska et al., 2018).

### Identification of novel CENP-A assembly factors

Proteins that we find uniquely involved in the CENP-A assembly process without any appreciable impact on CENP-A maintenance are the known dedicated CENP-A assembly factors Mis18α, Mis18β, M18BP1 and the CENP-A specific protein HJURP. Added to these we find CENP-R that was not previously implicated. Further we find ASF1B, a histone H3 chaperone involved in nucleosome recycling during DNA replication (Alabert and Groth, 2012). One hypothesis for its involvement in assembly may be that it acts as an acceptor protein for H3 exchange during CENP-A assembly. Further we find the **chromatin modifiers**, SMYD2 and NSD2^(WHSC1)^, (both SET domain-containing proteins) and SUV39H2, a methyltransferase for histone H3 lysine 9 tri-methylation (Rice et al., 2003), critical for heterochromatin function. The latter is of interest as heterochromatin has been implicated in CENP-A assembly and centromere formation (Folco et al., 2008; Olszak et al., 2011). The Polycomb proteins and histone methyl binding proteins CBX7 and L3MBTL (Chittock et al., 2017; Blackledge et al., 2015) involved in transcription repression are also found to significantly affect assembly of new CENP-A. Finally, we found several proteins involved in **ubiquitin metabolism**, UBE2A (Ubiquitin Conjugating Enzyme E2 A) a Rad6 homolog involved in H2B ubiquitylation (Kim et al., 2009), KEAP1 an adaptor protein for E3 ubiquitin ligase complexes and BRCC3 a BRCA1 and 2-associated Lys63-specific deubiquitinating enzyme (Feng et al., 2010), all affecting CENP-A maintenance.

Note that we discuss here the list of top candidates with an arbitrary cut-off of over 1.3 fold effect on CENP-A maintenance or 1.9 fold for CENP-A assembly. More genes were identified in the screen with a highly significant impact on CENP-A but with a lower fold difference (listed in supplemental tables S2, S3).

### A dynamic SUMO-cycle underlies the stable transmission of CENP-A chromatin

The candidate with the most significant impact on CENP-A maintenance as well as assembly is the SUMO protease SENP6. No other SUMO regulators were found having such high impact on CENP-A inheritance. SENP6 is involved in removal of SUMO2/3 chains from target proteins (Mukhopadhyay and Dasso, 2007). Loss of SENP6, both long-term and acutely, results in the loss of both ancestral and nascent CENP-A from chromatin. In fact, we find a rapid disassembly of the entire centromere and kinetochore complex and SENP6 is required for CENP-A chromatin integrity at any point in the cell cycle. These findings suggest that the centromere complex, including tightly bound chromatin components such as CENP-A are under continued surveillance by SUMO E3 ligases that control the localization of centromere proteins which is counteracted selectively by SENP6. A key future question is what is/are the specific target(s) of SENP6 within the centromere. CENP-C is a known SUMOylated centromere component that stabilizes CENP-A both *in vitro* and *in vivo* and is a central organizer of the centromere complex (Klare et al., 2015; Hendriks et al., 2017; Falk et al., 2015). SUMO control of CENP-C may therefore potentially play a role in maintaining CENP-A chromatin and thereby regulate the strength of the centromere.

In sum, we report the identification of a series of proteins not previously associated with centromere structure and function. These factors act selectively in the assembly of new CENP-A, in the maintenance of chromatin-bound CENP-A or both and reveal the dynamic nature of the maintenance of centromeric chromatin. The identification of these factors serves as resource for further discovery into the control of centromere assembly and inheritance.

## Supporting information

Table S1 List of screen siRNAs and genes

Table S2 CENP-A maintenance all candidates ranked

Table S3 CENP-A assembly all candidates ranked

## Acknowledgments

We thank Andrew Holland (Hopkins), Feng Zhang (MIT), Ronald Hay (Dundee), Song-Tao Liu (University of Toledo), Ian Cheeseman (Whitehead), Arshad Desai and Don Cleveland (both UCSD) for reagents. We thank Sebastiaan van den Berg for critically reading the manuscript. This work is supported by an ERC-consolidator grant ERC-2013-CoG-615638 and a Senior Wellcome Research Fellowship, both to LETJ. Further salary support to DLB was provided by Fundação para a Ciência e a Tecnologia (FCT) fellowship SFRH/BD/74284/2010 and an “Investigador FCT” position to LETJ.

## Author contributions

SM designed and performed the experiments, designed figures and drafted the paper. DLB and AFD performed initial pilot siRNA screen. JM provided tissue culture expertise and assisted in cell line construction. BN and SR provided essential guidance to high through put siRNA screening. CT build the data analysis pipeline. LET conceived the study, helped design the experiments, wrote the paper and acquired funding.

## Materials and methods

### DNA constructs

Constructs to build the SENP6^EGFP-AID/EGFP-AID^ cell line are as follows: The plasmid pX330-U6-Chimeric_BB-CBh-hSpCas9 from Feng Zhang lab [Addgene #42230, (Cong et al., 2013)] was used to construct the CRISPR/Cas vector plasmid according to the protocol as described in (Ran et al., 2013). Two guide RNA sequences: 5’ - GCAAGAGCGGCGGTAGCGCA - 3’ (sg1) and 5’ - GCCATGGATTAAGAAGGAGG - 3’ (sg2), designed to target the N terminal region of the SENP6 gene, were cloned into the pX330 backbone to generate the CRISPR/Cas vector plasmids pLJ869 (sg1) and pLJ870 (sg2) respectively. For generation of the N terminal AID tag, the construct LoxP-EGFP-LoxP-3xFLAG-miniAID-3xFLAG was gene synthesized and cloned into a pUc based vector to generate the template plasmid pLJ851. The homology donor vectors were constructed by PCR amplifying the template plasmid pLJ851 using Q5 DNA polymerase (New England Biolabs) with 110-base oligonucleotides using a 80-base homology sequence to the N terminal region of the SENP6 gene. The sequence of the upstream (US) and downstream (DS) homology arms are as follows: SENP6-US-HR-5’-CCGGCGCGGCCCCTCATCCCGGCGAGCACGGCGGCGGTGTGGGCCATGGATTAAGAAGGAGGCGGCGTGGGAGGAGGAAG’ and SENP6-DS-HR-5’-GCGGCCGGCAAGAGCGGCGGTAGCGCAGGGGAGATTACTTTTCTGGAAGGTACGTCTGTTTCTGCCCTTGACGGGGAGAAGGGAG’. In both cases homology arms were designed to introduce silent mutations in the PAM (protospacer-adjacent motif) recognition sequence after integration into the target locus in order to prevent Cas9 re-cutting.

### Cell lines and culturing conditions

All human cell lines were grown at 37°C, 5% CO2. Cells were grown in DMEM (Bio West) supplemented with 10% new born calf serum (NCS) (Bio West), 2 mM glutamine, 1 mM sodium pyruvate (SP) (Thermo Fisher Scientific), 100 U/ml penicillin and 100 µg/ml streptomycin. The SENP6 ^EGFP-AID/EGFP-AID^ cell line as shown in Figure 4A was constructed as follows: The parent cell line used was HeLa-CENP-A-SNAP clone #72 as described in (Jansen et al., 2007; Bodor et al., 2012). This cell line was transduced with pBABE-OsTir1-9Myc retrovirus (pLJ820) [gift from Andrew Holland, Johns Hopkins (Holland et al., 2012)] following the protocol as described in (Bodor et al., 2012). The infected cells were selected by 500 µg/ml of Neomycin (Gibco). Individual resistant cells were sorted by FACS. In order to generate genome targeted cell line, a single clone expressing OsTIR1 was amplified and grown in 10 cm dishes before transfecting them with CRISPR/Cas vector plasmids (pLJ869 and pLJ870) and homology donor derived from pLJ851 using Lipofectamine LTX (Invitrogen; Carlsbad, CA) according to manufacturer’s instructions. Monoclonal GFP positive clones were sorted by FACS. These clones were screened for homozygous tagging of the SENP6 gene by immunoblot using sheep anti-SENP6 antibody (gift from Ronald Hay, Dundee). Based on the immunoblotting results a single clone #18 was selected for performing auxin (Indole-3-acetic acid sodium salt or IAA; Sigma-Aldrich Cat. no. 15148) based experiments.. Auxin (IAA) was used at concentration of 500 µM.

### siRNA library

All siRNAs used in this study were obtained from Ambion Thermo Fisher Scientific as Silencer Select reagents. All siRNA sequences listed in Table S1 were blasted for unique target specificity against the current human genome using ENSEMBL V95, 2019 (EMBL, EBI and Wellcome Trust Sanger Institute).

### Transfection of siRNAs

All siRNAs used in this study were purchased from the Silencer Select collection of siRNAs from Thermo Scientific and are listed in Table S1. Production of RNAi microarrays as described in (Neumann et al., 2006). In brief, the siRNA-gelatin transfection solution was prepared in 384 well V-shaped plates using a manual 96 well pipetting device (Liquidator from Steinbrenner).For reverse solid transfection on cell arrays, 5 µl of 3µM siRNA was mixed with 1.75 µl of Lipofectamine 2000 (ThermoFisher), 1.75 µl H2O and 3 µl of Opti-MEM (ThermoFisher) containing 0.4 M sucrose and incubated for 20 min at room temperature (per well protocol). Next, 7.25 µl of 0.2% gelatin (w/v) was added and the mixture was printed on one-chamber Lab-Tek slides (Nunc) with a contact printer (BioRad) eight solid pins, giving a spot size of ∼400 µm diameter. The spot-to-spot distance was set to 1125 µm. Each Lab-Tek chamber accommodated 384 spots organized in 12 columns and 32 rows. The whole library of 2172 siRNAs was spotted onto seven Lab-Tek chambers, with 10 negative control siRNAs distributed randomly across each layout. After drying, the spotted library was seeded with CENP-A-SNAP pulse labeled cells as outlined in the section below.

For low throughput siRNA experiments, liquid-phase reverse transfection was performed on coverslips in 24 well plates using Lipofectamine RNAiMAX (Thermo Scientific) according to the manufacturer’s protocol with 3 × 10^4^ cells seeded per well. Cells were typically incubated for 48 hrs with the siRNA unless otherwise stated.

### SNAP Pulse-Chase and Quench-Chase-Pulse Labeling

Cell lines expressing CENP-A-SNAP were pulse labeled as previously described (Bodor et al., 2012). For the primary siRNA screen, HeLa-CENP-A-SNAP cells were grown in 6 well plates and pulse labeled with tetra-methyl-rhodamine-conjugated SNAP substrate (TMR-Star; New England Biolabs) at 4µM final concentration, labeling all pre-existing CENP-A molecules at the centromere. This was followed by a quenching step with bromothenylpteridine (BTP; New England Biolabs) at 2µM final concentration to prevent any further fluorescent labeling of nascent CENP-A following TMR-Star washout. The cells were then trypsinized (Trypsin-EDTA, Gibco) and 1.5 x10^5^ cells were directly seeded onto 1 chamber Lab-Tek slides spotted with the siRNA library as described above. The chambers were then incubated at 37°C, 5% CO2 for 48h. Finally, cells were labeled with Oregon-Green SNAP substrate (New England Biolabs) at 4 µM final concentration for labeling of the newly synthesized CENP-A molecules. Finally the cells were co-extracted (pre-extraction and fixation) in 4% paraformaldehyde, 0.2% Triton X and Hoeschst (1µg/ml) at room temperature for 30 min.

### Cell synchronization

Double thymidine-based synchronization was performed as described previously (Bodor et al., 2012). For G2 synchronization, cells released from double thymidine arrest were incubated 4 hrs later with CDK1 inhibitor RO-3306 (Merck millipore) at 9µM concentration for 4 hrs. For mitotic arrest and release cells were incubated with 2.5 µM of EG5 inhibitor Dimethylenastron for 13h. Following inhibitor washout, cells were released for 9 hrs to obtain a synchronous population of late G1 cells.

### Immunofluorescence procedures

The immunofluorescence procedures were followed as described (Bodor et al., 2012). Briefly, the cells were grown on glass coverslips coated with poly-L lysine (Sigma-Aldrich) and fixed with 4% formaldehyde (Thermo Scientific) for 10 min followed by permeabilization in PBS with 0.1% Triton-X-100. When staining with antibodies for CENP-A, CENP-C, CENP-H, DSN1, NNF1 and HEC1, an additional pre-extraction step was included which involved incubation of cells with 0.1% Triton-X-100 in PBS for 5 min prior to fixation by formaldehyde. The following antibodies and dilutions were used: mouse monoclonal anti-CENP-A [gift from Kinya Yoda (Nagoya University) (Ando et al., 2002)] at 1:100; rabbit polyclonal anti-CENP-B (sc22788; Santa Cruz Biotechnology, Dallas, TX) at 1:100; mouse monoclonal anti-CENP-B isolated from hybridoma line (2D-7) (Earnshaw et al., 1987) at 1:50; mouse monoclonal anti-CENP-C isolated from hybridoma line (LX191) [gift from Don Cleveland, UCSD (Foltz et al., 2009)] at 1:100, rabbit polyclonal anti CENP-T [gift from Don Cleveland, UCSD (described in (Barnhart et al., 2011); rat monoclonal anti-CENP-H and rabbit polyclonal anti-CENP-I (both gifts from Song-Tao Liu, University of Toled0); rabbit polyclonal anti-DSN1 (gift from Iain Cheeseman, Whitehead) at 1:100; rabbit polyclonal anti-NNF1 (gift from Arshad Desai, UCSD); mouse monoclonal anti-HEC1 (Thermo Scientific Pierce MA1-23308), sheep polyclonal anti-SENP6 (gift from Ronald Hay, Dundee) at 1:500; rat monoclonal anti-Tubulin (SC-53029, Santa Cruz Biotechnology, Dallas, TX) at 1:10,000 and mouse monoclonal anti Cyclin-B (SC-245, Santa Cruz Biotechnology, Dallas, TX) at 1:50. All primary antibody incubations were performed at 37°C for 1h in a humid chamber. Fluorescent secondary antibodies were obtained from Jackson ImmunoResearch (West Grove, PA) or Rockland ImmunoChemicals and used at a dilution of 1:200. All secondary antibody incubations were performed at 37°C for 45 min in a humid chamber. Cells were stained with DAPI (4’,6-diamidino-2-phenylindole; Sigma-Aldrich) before mounting in Mowiol. EdU (5-ethynyl-2’-deoxyuridine) labelling was performed for 15 min as per the manufacturer’s instructions (C10340, Life Technologies) in order to stain S phase nuclei. In the experiments where EdU labeling was performed, EGFP-SENP6 signal was detected using GFP-Booster Atto488 (Chromotek).

Immunofluorescent signals of Figures 2, 3, 4 and S3 were quantified using the CRaQ (Centromere Recognition and Quantification) method as described previously (Bodor et al., 2012) using CENP-B as centromeric reference. Hec1, Dsn1 and Nnf1 levels were measured only in prometaphase or metaphase (based on DAPI staining) nuclei. The immunofluorescent signal of EGFP-SENP6 in Figure S2 was measured using an ImageJ based macro which measured the median intensity of the whole nucleus.

### Microscopy

Imaging for the primary siRNA screen was performed on an Olympus ScanR (IX-81) automated inverted microscope (Olympus Biosystems), equipped with a Hammamatsu Orca-ER and a MT20 light source. The microscope was controlled by ScanR acquisition software (version 2.3.0.7). A 20x 0.75 NA air objective (UpLANsaPO; Olympus Biosystems) was used for the primary screening on cell arrays and a single plane image was acquired for each siRNA spot. Filter settings and exposure times were the following: Dapi: Ex: 347/50 EM: 460/50, exposure time 20ms; Oregon Green Ex: 482/35 EM: 536/40, exposure time 500 ms and TMR star: 545/30 EM: 610/72 exposure time 1500 ms.

For validation and characterization experiments imaging was performed using a Deltavision Core system (Applied Precision) inverted microscope (Olympus, IX-71) coupled to Cascade2 EMCCD camera (Photometrics). Images (512 x 512) were acquired at 1X binning using a 100x oil objective (NA 1.40, UPlanSApo) or a 60x oil objective (NA 1.42 PlanApoN) with 0.2 µm z sections.

### Image Analysis

Image analysis of the primary screen was performed using a CellProfiler (Carpenter et al., 2006) pipeline, available for download (https://git.embl.de/grp-almf/sreyoshi-mitra-jensen-centromere-screen/blob/master/publication/mitra_centromere_screen_cp2.2.0.cpproj) and can be used with CellProfiler version 2.2.0. In brief, nuclei were detected in the DAPI image using automated thresholding. Based on their texture, non-interphase (mitotic or other, e.g. dead cells) nuclei were removed from the analysis. The ‘old’ and ‘new’ CENP-A-SNAP images were subjected to a morphological tophat filter in order to remove diffuse (non-centromeric) background signal. Centromeric regions were detected using automated thresholding and, for each nucleus, the integrated intensity within centromeric regions was measured. For downstream data analysis, we computed the mean values of all nuclei in each image, yielding, for each image, two measurements: “Mean_Interphase Nuclei_Intensity_Integrated Intensity_New Tophat Centromere Mask” and “Mean_Interphase Nuclei_Intensity_Integrated Intensity_Old Tophat Centromere Mask”. In addition, to be able to reject out-of-focus images, we used CellProfiler’s Measure Image Quality module, specifically the values: ImageQuality_Power Log Log Slope_Nucleus and ImageQuality_Power Log Log Slope_Old. Those are spatial frequency based measurements, where low values indicate missing high spatial frequencies such as it is the case for out-of-focus images.

### Statistical Analysis

Visual inspection, quality control and statistical analysis of the primary screen was performed using HTM explorer (https://github.com/tischi/HTM_Explorer). Primary data was the CellProfiler output table, containing measurements of 13440 images, corresponding to 35 384 spotted Lab-Tek chamber. In terms of quality control, we filtered out-of-focus images, rejecting all images where either ImageQuality_PowerLogLogSlope_Nucleus < −2.3 or ImageQuality_PowerLogLogSlope_Old < −2.0, thereby removing 596 images from the analysis. The threshold values −2.3 and −2.0 were determined by visual inspection. The aim of the primary screen was to measure the siRNA knockdown induced fold-change of CENP-A centromere signal relative to our negative control siRNA. To this end, we first computed a log2 transform of both CENP-A readouts (Mean_InterphaseNuclei_Intensity_IntegratedIntensity_NewTophatCentromereMask and Mean_InterphaseNuclei_Intensity_IntegratedIntensity_OldTophatCentromereMask). Next, for both readouts, we performed a Lab-Tek chamber-wise normalization by subtracting, for each chamber, the mean value of the 10 negative control images on that plate. To obtain one final score per treatment (siRNA) we pooled the normalized values for each treatment across all plates and performed a t-test against the normalized negative control values from each plate. The t-test’s estimate of the difference between treatment and control represents the log2 fold ratio of treatment and control. The t-test’s p-value represents the statistical significance of this difference.

### Parameters as listed in the data tables (Supplemental tables S2 and S3)

Fold difference vs control (t-test position estimate): In the first step, data to control measurements from a single chamber is normalized by subtracting the mean of the control positions in that chamber. Afterwards, for one specific treatment all chambers are identified that contained this treatment and all treatment and control measurements from these batches are pooled. The t test positions estimate is the difference of treated positions and control positions (after batch correction). If the log2 data transformation of this value is chosen, this difference gives the fold-change of treatment vs. control (in log2 scale). To compute the actual fold change you can use this formula: 2^estimate^

Significance (t-test p value): After performing batch correction, for one specific treatment all chambers are identified that contained this treatment and all treatment and control measurements from these batches are pooled; a t-test is performed of the treated positions against the control positions. t test positions p value gives the p value computed from the above t test.

Median z score: For each chamber, the data of all images within one position are averaged such that we have one number per position. Then a z-score is computed for each position as Z = (value - mean (ctrls)) / sd(ctrls) Where the mean and standard deviation (sd) are computed across all positions that contain the selected control measurements. Median z-score is the median value of z-scores from multiple chambers containing the specific treatment.

Median robust z score: Same as median z score, but for each chamber a z-score is computed as Z = (treated - median (ctrls)) / mad(ctrls), where mad is the so called median average deviation (a median based analog to the standard deviation).

To represent the final dataset in Figure 1, the Fold difference vs control [t test position estimate values (equivalent to log_2_ fold change of siRNA treatment versus control)] and the negative log_10_ *p*-values were plotted as volcano plots using GraphPad Prism version 7. Two volcano plots were generated depicting the candidates affecting the loading of new CENP-A molecules and those affecting the maintenance of pre-assembled old CENP-A at the centromere, hereafter called ‘loading’ and ‘maintenance’ candidates respectively. For the maintenance candidates, a cut-off of −0.4 was set for log_2_ fold-change representing at least 1.3-fold reduction in old CENP-A intensity in the siRNA treated condition versus that in a negative scrambled siRNA control. For the loading candidates, a cut-off of −1 was set for log_2_ fold-change representing at least 2-fold reduction in new CENP-A intensity in the siRNA treated condition versus that in a negative scrambled siRNA control. A cut-off value of 3 for –log (*P*-value), corresponding to a *P*-value of <0.001, was employed to ascribe statistical significance for both maintenance and loading candidates.

### Immunoblotting

Whole cell extracts were prepared by direct lysis in 1X Laemmli sample buffer, separated by SDS-PAGE and transferred onto nitrocellulose membranes. The following antibodies and dilutions were used: rabbit polyclonal anti-CENP-A (#2186, Cell Signaling Technology) at 1:500; mouse monoclonal anti-α Tubulin (T9026, Sigma-Aldrich) at 1:5000; rabbit polyclonal anti-CENP-C (Covance) gift from Don Cleveland, UCSD) crude serum at 1:5000 and sheep polyclonal anti-SENP6 (gift from Ronald Hay, Dundee) at 1:5000. IRDye800CW-coupled anti-rabbit (Licor Biosciences), DyLight800-coupled anti-rabbit (Rockland Immunochemicals) and DyLight680-coupled anti-mouse (Rockland Immunochemicals) secondary antibodies were used at 1:10,000 prior to detection on an Odyssey near-infrared scanner (Licor Biosciences).

## Supplemental figures

**Supplemental Figure S1. (related to Figure 1).**
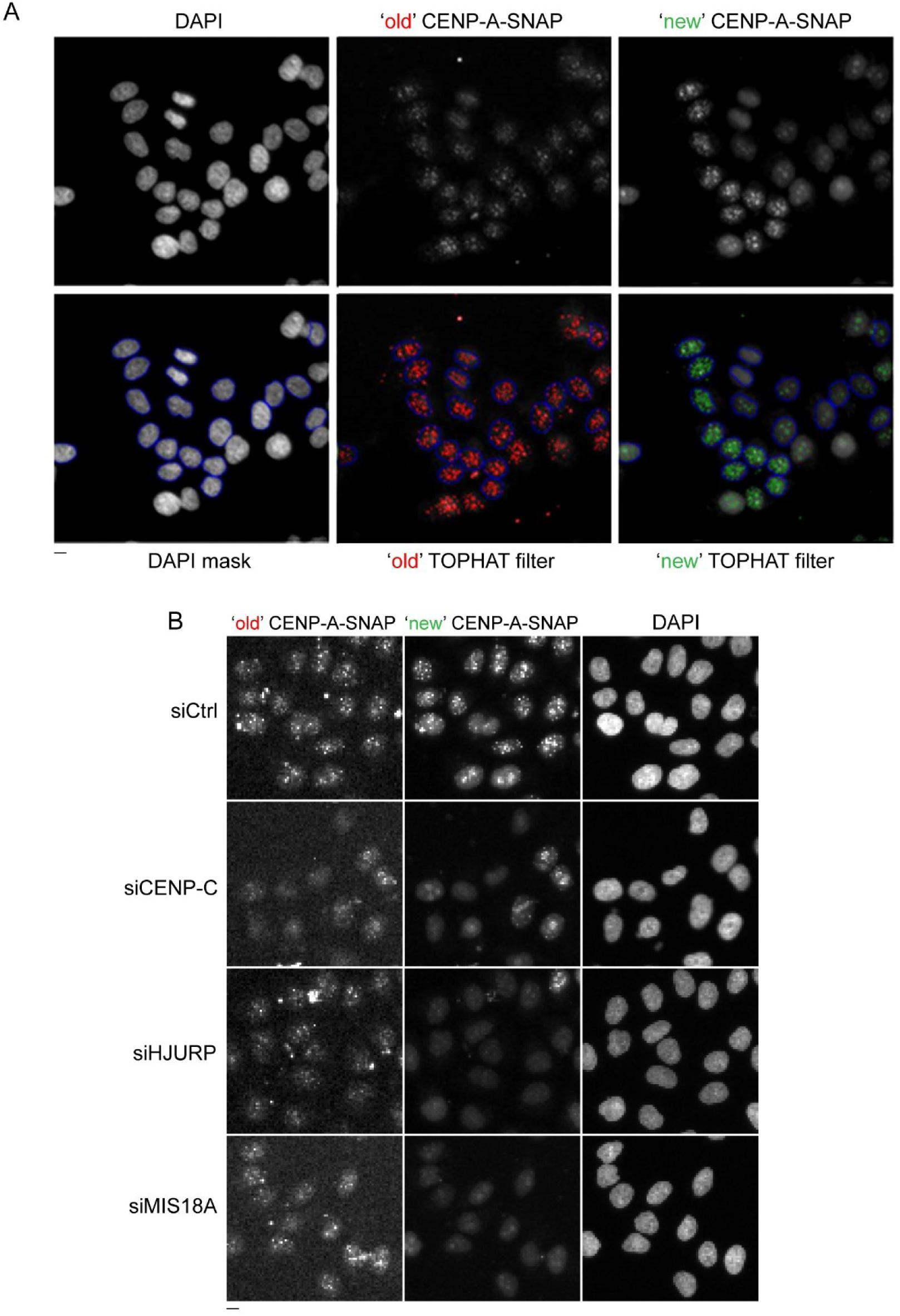
Quantification of siRNA screen results by Cell Profiler pipeline. **(A)** Representative low magnification images depicting the various steps of the custom Cell Profiler pipeline used to quantify ‘old’ and ‘new’ CENP-SNAP fluorescent signals in the primary siRNA screen. A DAPI mask was applied in order to identify interphase nuclei. Within the DAPI mask, custom TOPHAT filters were applied based on signal area and intensity in order to identify centromeres and measure ‘old’ and ‘new’ CENP-A-SNAP signal intensities. **(B)** Representative images for ‘old’ (siCENP-C) and ‘new’ (siHJURP, siMIS18A) CENP-A-SNAP phenotypes used as control siRNAs in the screen.

**Supplemental Figure S2. (related to Figure 4).**
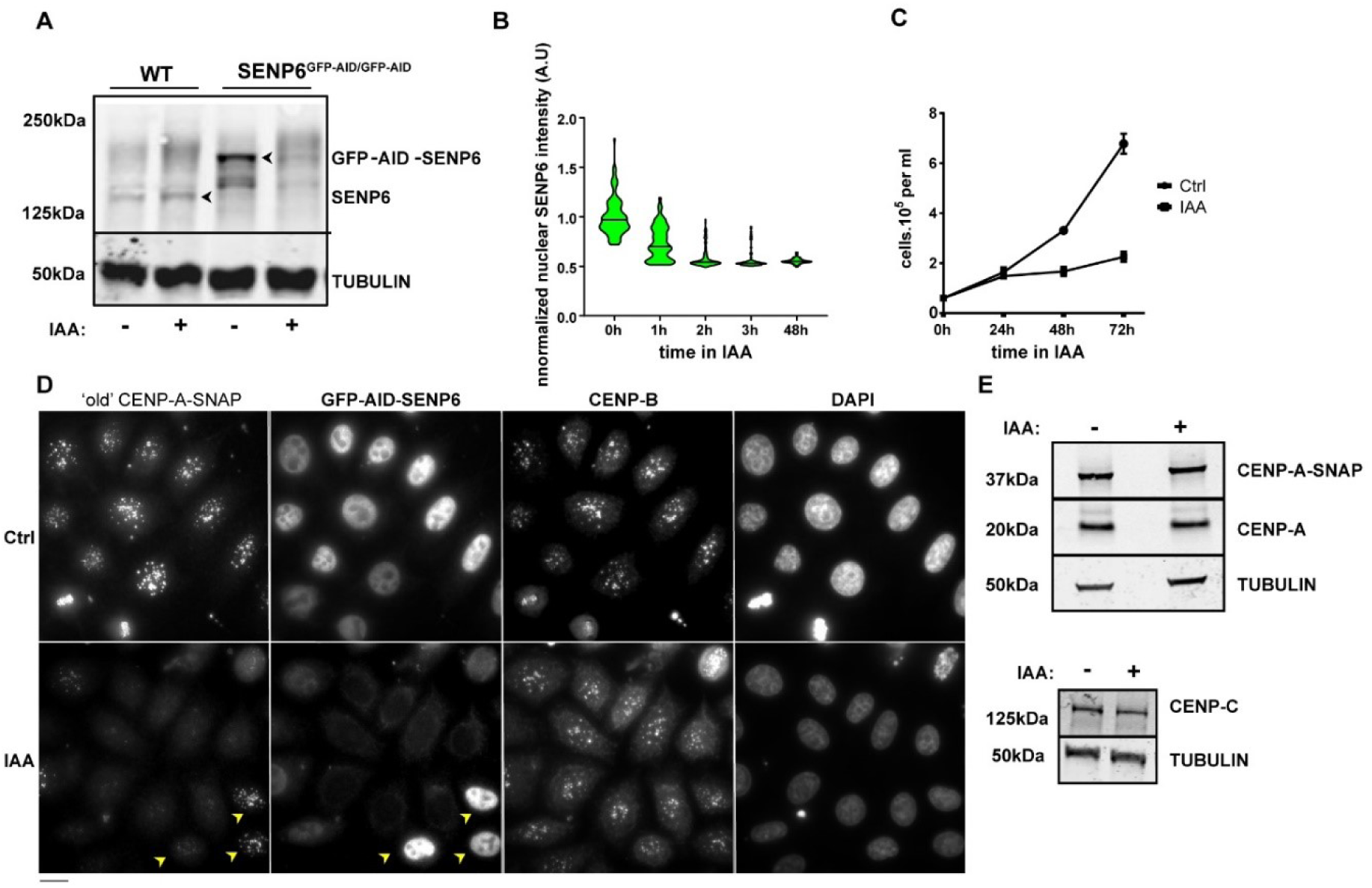
Auxin (IAA) treatment leads to rapid degradation of AID-tagged SENP6 and subsequent loss of centromeric CENP-A and cell growth arrest. **(A)** Western blot showing the degradation of GFP-AID-SENP6 protein upon addition of auxin (IAA) in a homozygously tagged (SENP6 ^GFP-AID/GFP-AID^) cell line. Untagged wild-type (WT) and SENP6 ^GFP-AID/GFP-AID^ cell lines were treated with auxin or mock-treated for 24 hours, extracts were separated by SDS-PAGE and immunoblotted with anti-SENP6 antibody. Tubulin was used as the loading control. Arrowheads indicate the WT and tagged SENP6 respectively. **(B)** Kinetics of auxin (IAA)-mediated degradation of GFP-AID-SENP6 measured by fluorescence microscopy and normalized to the mean intensity at 0 hour time point is plotted as violins. At least 200 nuclei were measured under each condition. Bar indicates the median value. **(C)** Long term auxin (IAA)-mediated depletion of SENP6 leads to cell division arrest. Cell numbers were measured as a function of time under auxin (IAA) treatment or mock-treatment (Ctrl). Two replicate experiments were performed. Bars indicate SEM. **(D)** Images of centromeric levels of pre-incorporated ‘old’ CENP-A-SNAP in a pulse-chase assay following auxin (IAA)-mediated depletion of GFP-AID-SENP6 for 24 hours as quantified in Figure 4C. Cells were counterstained with CENP-B as centromeric reference. Yellow arrowheads indicate nuclei which retain GFP-AID-SENP6 after auxin (IAA) treatment and correspondingly retain ‘old’ CENP-A-SNAP. Bar, 10 µm. **(E)** Western blots showing the total levels of CENP-A and CENP-C proteins after degradation of GFP-AID-SENP6 by auxin (IAA) treatment for 24 hours. Extracts from auxin (IAA) treated (+) or control (−) cells were separated by SDS-PAGE and immunoblotted with anti-CENP-A (detecting both SNAP tagged and endogenous CENP-A) or anti-CENP-C antibody. Tubulin was used as loading control.

**Supplemental Figure S3. (related to Figure 4).**
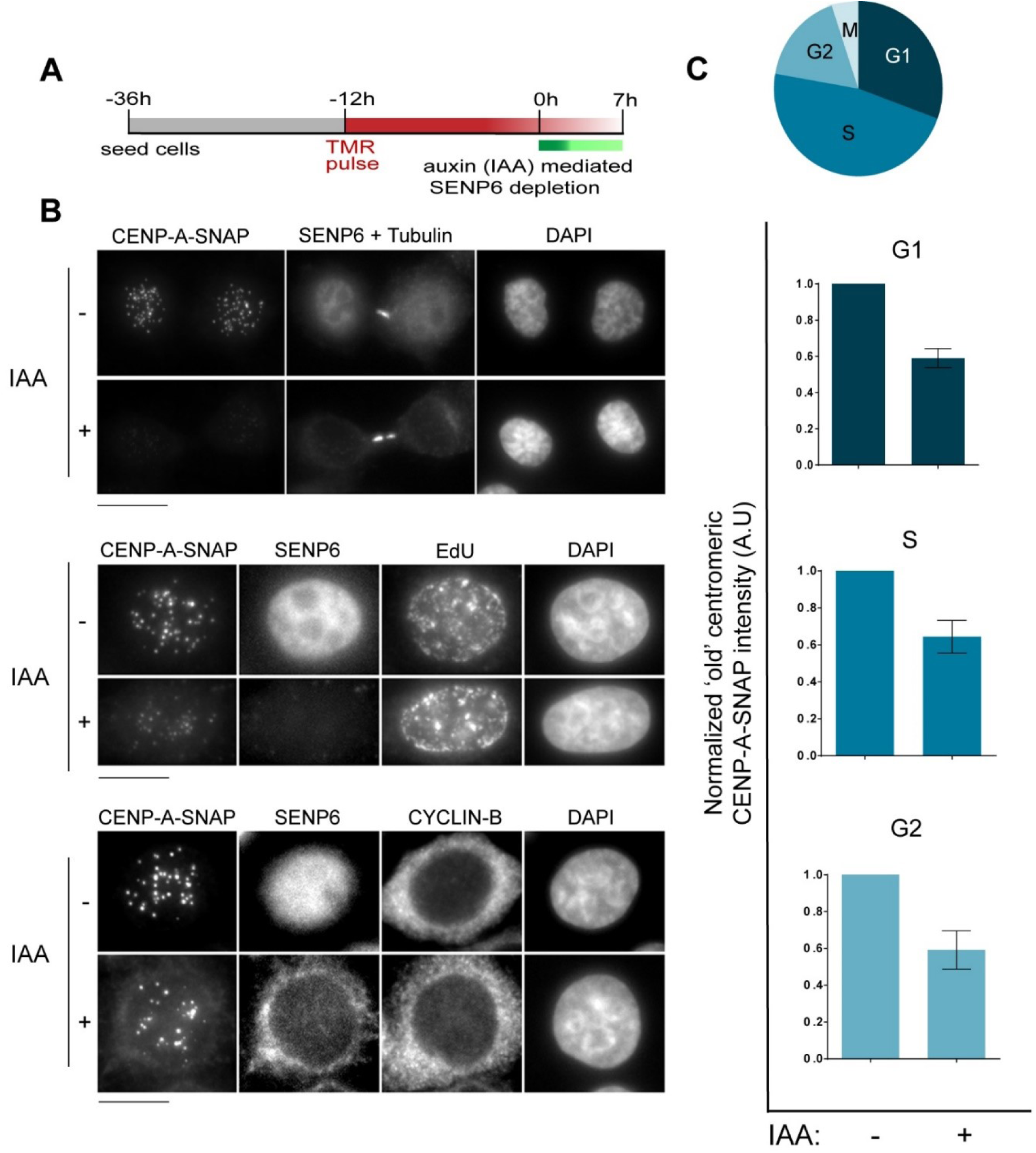
Auxin (IAA) mediated degradation of SENP6 in unsynchronized cells shows its requirement for maintaining centromeric CENP-A at all cell cycle stages. **(A)** Experimental scheme of CENP-A-SNAP pulse-chase assay following auxin (IAA) mediated depletion of SENP6 in unsynchronized cells. **(B)** Results of the experiment described in (A) showing images of centromeric levels of ‘old’ CENP-A-SNAP following auxin (IAA) mediated depletion of SENP6 at specific cell cycle stages. Cells were counter-stained with α-tubulin to score for mid-body positive G1 cells, EdU to score for S phase cells and with Cyclin B to stain G2 cells. **(C)** (Top panel) Schematic depicting relevant stages of the cell cycle. (Bottom panel) Quantification of (B). Centromeric CENP-A-SNAP signal intensities in cells under auxin (IAA) treatment (+) were normalized to the control mock-treated condition (−) in each experiment and plotted along the y-axis. Three replicate experiments were performed. Bars indicate SEM.

**Supplemental table 1.** List of all 2172 siRNA sequences with their respective gene targets and ENSEMBL IDs

**Supplemental table 2.** Ranking of all siRNA targets according to their impact on CENP-A maintenance in SNAP pulse-chase screen. Ranking is based on Fold difference “t test positions estimates”. Scores for Significance “t test positions p value” with respective sign code, Z scores and number of objects (cell nuclei) are also listed. The descriptors are defined in methods.

**Supplemental table 3.** Ranking of all siRNA targets according to their impact on CENP-A assembly in SNAP quench-chase-pulse screen. Ranking parameters are as in Supplemental table S2 and are defined in the methods.

